# Quantitative relationships between SMAD dynamics and target gene activation kinetics in single live cells

**DOI:** 10.1101/491894

**Authors:** Onur Tidin, Elias T. Friman, Felix Naef, David M. Suter

## Abstract

The transduction of extracellular signals through signaling pathways that culminate in a transcriptional response is central to many biological processes. However, quantitative relationships between activities of signaling pathway components and transcriptional output of target genes remain poorly explored. Here we developed a dual bioluminescence imaging strategy allowing simultaneous monitoring of nuclear translocation of the SMAD4 and SMAD2 transcriptional activators upon TGF-β stimulation, and the transcriptional response of the endogenous *connective tissue growth factor (ctgf)* gene. Using cell lines allowing to vary exogenous SMAD4/2 expression levels, we performed quantitative measurements of the temporal profiles of SMAD4/2 translocation and *ctgf* transcription kinetics in hundreds of individual cells at high temporal resolution. We found that while nuclear translocation efficiency had little impact on initial *ctgf* transcriptional activation, high total cellular SMAD4 but not SMAD2 levels increased the probability of cells to exhibit a sustained *ctgf* transcriptional response. The approach we present here allows time-resolved single cell quantification of transcription factor dynamics and transcriptional responses and thereby sheds light on the quantitative relationship between SMADs and target gene responses.

## Introduction

Cells relay information from environmental stimuli through signaling pathways to modulate gene expression. Over the past decade, numerous studies have shed light on the dynamics of transcription factor shuttling and the resulting transcriptional and translational outputs in response to extracellular signaling ^1–3^. The transcriptional response to extracellular stimuli has been shown to exhibit surprisingly large variability among phenotypically identical individual cells. This variability stems not only from stochasticity inherent to biochemical processes ^4^, but also from variations in the expression level or state of a large number of factors involved in signaling pathway transduction or gene expression components ^5^. However, how the variability in expression level or activity of upstream components is quantitatively related to variability in the transcriptional response of target genes is poorly understood. More recently, methods allowing to measure multiple nodes in signaling pathways were developed and applied successfully to study several pathways in live cells ^6–9^, but simultaneous monitoring of transcription factor activity and transcriptional kinetics of endogenous target genes remains challenging.

The TGF-β superfamily signaling pathway plays a central role in a broad range of biological processes, such as embryonic development, tissue homeostasis and cancer ^10,11^. The pathway has two main branches activated at the transmembrane receptor level by specific binding of ligands in the TGF-β superfamily. Among those ligands, TGF-β signals through a transmembrane receptor that recruits SMAD2/3 and allows their phosphorylation. pSMAD2/3 subsequently heterodimerizes with SMAD4 to translocate into the nucleus and activate hundreds of target genes in different cellular contexts ^12–14^. Single-cell studies have revealed the pulsatile nature of SMAD shuttling dynamics and the heterogeneity of signaling determined by varying protein levels of individual cells ^15,16^. Yet, how cells interpret SMAD signaling and elicit a response remains unclear, mostly due to the scarcity of experimental systems allowing simultaneous measurements of SMAD dynamics and transcriptional output in the same cells. One study decoded the contributions of SMAD dynamics to downstream response in the TGF-β pathway using synthetic TGF-β inducible reporters, and demonstrated how the kinetics of ligand presentation impacts SMAD translocation and its target gene response ^17^. However, that study did not investigate SMAD translocation activity and target gene response in the same cells, and used a synthetic TGF-β targeted promoter construct, which may differ in its response as compared to an endogenous TGF-β target gene. Similarly, another study investigated SMAD-mediated target gene transcriptional activity and revealed that cells interpret fold-changes rather than absolute concentrations of TGF-β to elicit downstream responses ^18^. However, in that study, target gene response analysis relied on analysis of fixed cells by single-molecule FISH (sm-FISH), which does not allow to capture the full range of information on response dynamics; moreover, the long-term dynamics of the target gene response was not explored.

Among the direct targets of the TGF-β signaling pathways, *connective tissue growth factor* (*ctgf*) encodes a secreted factor that promotes fibroblast proliferation and fibrosis, and plays a central role during wound repair as well as numerous pathological fibrotic conditions ^19^. Using gene trapping of a short-lived luciferase reporter, we have previously shown that *ctgf* (similarly to most mammalian genes) is transcribed in a temporally discontinuous manner referred as to transcriptional bursting ^20^. We have further shown that TGF-β stimulates *ctgf* transcription by increasing the transcription rate of *ctgf* during transcriptionally active temporal windows ^21^. However, the quantitative relationships between components of the TGF-β signaling pathways and the transcriptional output of *ctgf* are poorly understood.

Here we aimed at understanding how SMAD4 and SMAD2 nuclear translocation dynamics and expression levels quantitatively relate to *ctgf* transcriptional activity. We generated cell lines allowing to modulate SMAD4 and SMAD2 expression levels, and to simultaneously monitor their nucleo-cytoplasmic shuttling and the transcriptional activity of *ctgf* by two-color live luminescence imaging of single cells. We found that while the increase of SMAD4 and SMAD2 nucleo/cytoplasmic ratio were poor predictors of the transcriptional response of *ctgf*, high SMAD4 but not SMAD2 expression increased the probability of exhibiting sustained ctgf transcriptional activity upon TGF-β stimulation.

## Results

### Simultaneous monitoring of transcription factor shuttling and target gene activation in single living cells

We previously generated an NIH-3T3 mouse fibroblast gene trap cell line expressing a short-lived luciferase protein allowing to monitor transcriptional activity of the *ctgf* gene in single live cells by luminescence microscopy (GT:ctgf) ^20^. To allow live monitoring of SMAD4 and SMAD2 nucleo-cytoplasmic shuttling in the same cells, we established two doxycycline (dox)-inducible stable cell lines each expressing a fusion protein of a luminescence (Nanoluciferase (Nluc), ^22^ reporter to either SMAD4 or SMAD2 (Fig.1A and 1B), referred as to iS4 and iS2 cells lines, respectively. We reasoned that low-level expression of these exogenous fusion proteins should allow monitoring nucleo-cytoplasmic shuttling of the SMADs without significantly altering the total pool of SMAD4/2. To determine the optimal dox concentration, we first characterized the dox dose-response in the expression of SMAD4/2-Nluc (Fig.1C-F), and found that 2ng/ml of dox treatment allowed expression levels lower or in the same range as endogenous SMAD4/2 (Fig.1D and 1F). We also monitored SMAD4 nuclear import in the GT:ctgf cell line and SMAD2 phosphorylation in the iS2 cell line treated with 2ng/ml of dox after TGF-β stimulation. As expected, SMAD4 was rapidly shuttled to the nucleus and SMAD2 phosphorylation reflected the response dynamics to TGF-β stimulation.

We then performed time-lapse, two-color luminescence imaging of either SMAD4/2-Nluc with Fluc driven by the endogenous *ctgf* regulatory sequences at a temporal resolution of 5 minutes. Since luminescence imaging does not involve sample illumination, it does not suffer from photobleaching or phototoxicity, thus allowing to image cells for long periods of time (up to several days) with high sensitivity and at high temporal resolution. While the substrate for Nluc is unstable in the medium and thus Nluc signal decreased over long timescales, this did not impact our ability to quantify SMAD4/2 nucleo-cytoplasmic shuttling as this is a ratiometric measurement ^23^. In contrast to unstimulated cells or cells treated with the TGF-β receptor antagonist SB-431542 (Supplementary Figures 1C-D), we observed robust nuclear shuttling of SMAD4 and SMAD2 (Fig.1G and Supplementary Figures 1E-F) and the subsequent transcriptional response of *ctgf* (Fig.1H, Supplementary Figure 1E and Supplementary Figure 1G) upon TGF-β stimulation. Therefore, dual-color luminescence imaging allows simultaneous recording of transcription factor shuttling and transcriptional responses at high temporal resolution.

**Figure 1.**
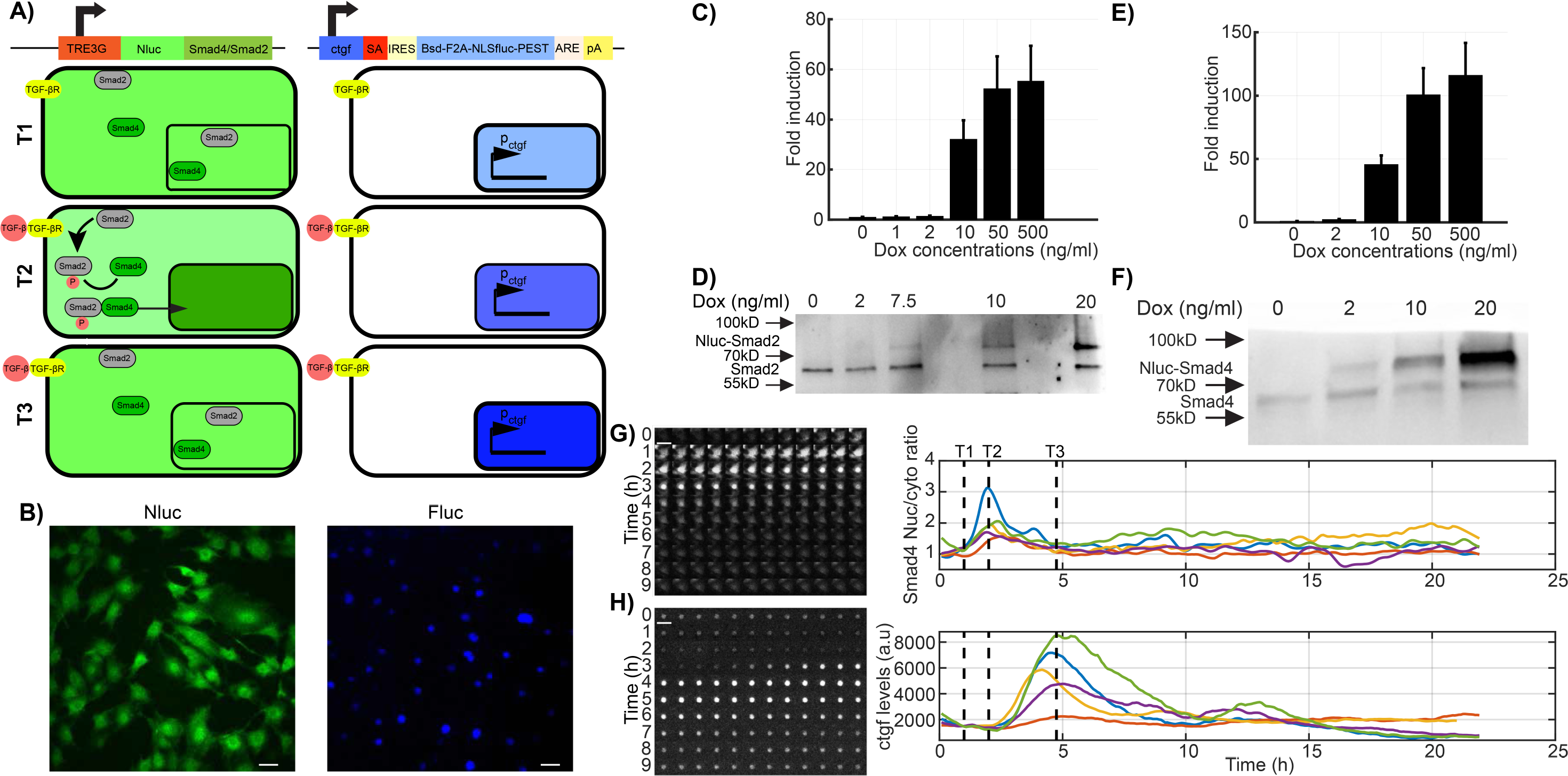
Dual bioluminescence reporter system. A) Schematic illustration of the dual bioluminescent reporter system. Nanoluciferase (Nluc) was fused to Smad4 and expressed under the control of the TRE3G doxycycline promoter (left). A short-lived NLS-luciferase is integrated into the endogenous *ctgf* locus (right, ^20^). Bsd: Blasticidin-deaminase; F2A: foot-and-mouth virus co-translationally cleaved peptide; NLS: Nuclear localization signal; Fluc: firefly luciferase; PEST: protein destabilizing sequence. In unstimulated condition, SMAD4 is localized both in the nucleus and cytoplasm (T1) and *ctgf* is expressed at a basal level. TGF-β stimulation results in a transient nuclear enrichment of SMAD4 signal while the Fluc signal increases. T2 and T3 represent the time points after TGF-β stimulation at which the average nuclear SMAD localization and the *ctgf* response reach their maxima, respectively. B) Dual bioluminescence detection of NLuc-SMAD4 and ctgf levels (Fluc) in NIH-3T3 cells. Image shows one frame obtained from luminescence movies with Nluc and Fluc signals shown in green and blue, respectively. C) Fold-change of Nluc-SMAD2 expression (luminescence) with different doses of doxycycline. D) Western blotting analysis of Nluc-SMAD2 and SMAD2 expression with different doses of doxycycline. The cropped blot is used in the figure and the full length Western blot scans for the cropped images are shown in Supplementary Figure 5. E) Fold-change of Nluc-SMAD4 expression (luminescence) with different doses of doxycycline. F) Western blotting analysis of Nluc-SMAD4 and SMAD4 expression with different doses of doxycycline. The cropped blot is used in the figure and the full length Western blot scans for the cropped images are shown in Supplementary Figure 5. G-H) Time series images of a tracked single cell expressing Nluc-SMAD4 on both Nluc (G) and Fluc (H) channel in the luminescence microscopy. Time goes from top left to bottom right, with a time resolution of 5 minutes. Single-cell quantification of SMAD4 translocation (upper right panel) and ctgf expression level (lower right panel) profiles in five individual cells stimulated with TGF-β (5nM) at time T1. Scale bar: 20 μm. Error bars: mean ± SD; n = 3.

### Quantitative relationship between SMAD nuclear import dynamics and *ctgf* response

We next monitored SMADs nuclear import and *ctgf* responses in hundreds of individual cells, in both iS4 and iS2 cell lines treated with 2ng/ml of dox and stimulated with 5nM of TGF-β. We found that almost all cells rapidly increased nuclear SMAD4 after stimulation, reaching a peak 1h after stimulus (Fig.2A-B). An increase in transcriptional activity of *ctgf* was observed on average 7 minutes after SMAD4 reached its maximal nuclear concentration (Fig.2E, green and blue dashed lines). Individual cells displayed little variability in the timing of SMAD4 translocation (Fig.2F) but larger variability in its nucleo/cytoplasmic ratio (Fig.2G), suggesting variable transduction efficiency of TGF-β signaling to SMAD4 shuttling. Both timing and amplitude of the *ctgf* transcriptional response displayed a broad distribution, reflecting large cell-to-cell variability in upregulation of the *ctgf* gene (Fig.2F and 2H). We also performed the same experiments on the iS2 cell line and observed similar translocation dynamics and gene expression response profiles (Fig.2I-P). We conclude that SMAD4 and SMAD2 translocation timings are tightly controlled, but translocation efficiencies and the transcriptional responses of *ctgf* varied over a ~ 3-fold range. Since both the translocation efficiencies of the SMADs and *ctgf* responses were broadly distributed, we next aimed at determining whether individual cells displayed correlated SMADs translocation and *ctgf* response profiles. Surprisingly, the SMAD4 and SMAD2 translocation amplitudes were not correlated to the amplitude of the *ctgf* response (Fig.3A-B), suggesting that upon treatment with 5nM of TGF-β, SMAD4/2 nuclear import is not limiting for transcriptional activation of *ctgf*. We then verified whether SMAD4/2 become limiting using lower doses of TGF-β (Supplementary Figure 2). At a concentration of 500, 50 and 5 pM, the amplitudes of SMAD4/2 nuclear import were again not significantly correlated to the *ctgf* response (Fig.3C-H), suggesting that SMAD4/2 shuttling is generally not rate-limiting in the transcriptional response of *ctgf* in our system. In contrast, the timings of SMAD4 and SMAD2 import peak were weakly, but significantly positively correlated to the initiation of the *ctgf* response (Fig.3I-J). This suggests that variability in the timing of the TGF-β signaling upstream of SMAD4 is propagated to transcriptional activation of *ctgf*.

**Figure 2.**
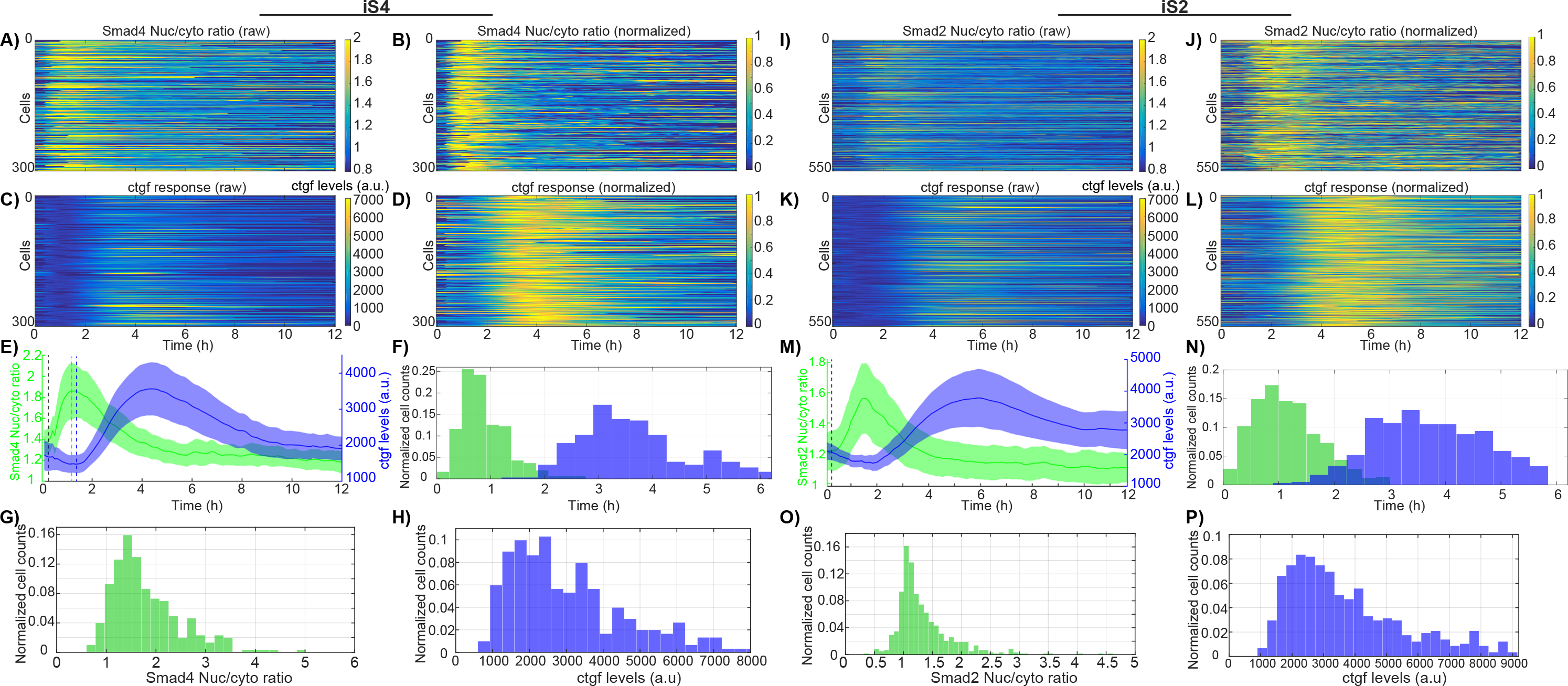
Single cell variability of SMAD4/2 translocation dynamics and *ctgf* expression. A-D) and I-L) Heatmaps of single cell SMAD4/2 translocation (A-B and I-J) and *ctgf* expression (C-D and K-L) upon 5nM TGF-β stimulation. Data is represented both in absolute luminescence levels (A,C and I,K) and intensities normalized on the maximal intensity for each trace (B,D and J,L). E) and M) Population average of SMAD4/2 translocation and *ctgf* expression. F and N) Single cell distribution of peak timing for SMAD4/2 translocation (green) and *ctgf* expression (blue). SMAD4: Mean = 55 min, CV=0.33; SMAD2: Mean = 66 min, CV= 0.35. G and O) Single-cell distribution of nuclear to cytoplasmic ratio at translocation peak for SMAD4 (G) and SMAD2 (O). H and P) Single-cell distribution of *ctgf* expression peak levels for iS4 (H) and iS2 (P) cell lines.

**Figure 3.**
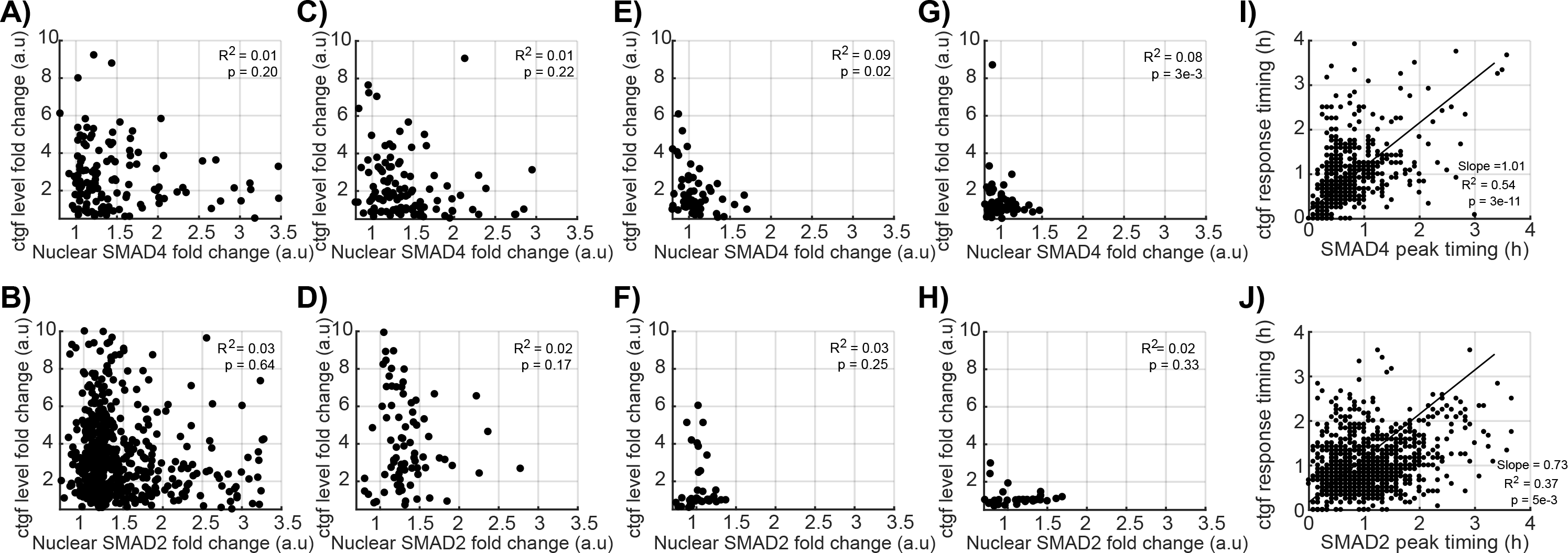
Quantitative single-cell relationship between SMAD4/2 translocation and ctgf expression. A-H) Correlation between the fold-change in SMAD4 (A, C, E) and SMAD2 (B, D, F) nuclear localization with ctgf response, after stimulation with 5nM (A and B), 0.5nM (C and D), 0.05nM (E and F), 0.005nM (G and H) of TGF-β. I-J) Correlation between the SMAD4 (I) and SMAD2 (J) translocation peak time and the time delay before *ctgf* expression starts to rise.

### Analog encoding of TGF-β concentration information by dose-dependent SMAD signaling and *ctgf* responses

While some early studies assume that TGF-β concentrations stay constant over time after its addition, it has been shown that TGF-β is internalized and degraded by cells and is determinant for the downstream signaling ^24,25^. To determine whether the ligand dose-response is analog (graded response of all cells) or digital (modulation of the fraction of responding cells) we performed stimulation experiments with a range of TGF-β doses (5pM-5nM) and quantified *ctgf* responses together with SMAD4/2 profiles in single-cells. These experiments revealed a dose-dependent (analog) profile of the response characterized by gradually altered SMAD4/2translocation and the target gene activity (Fig.4A-B), consistent with previous reports ^16^. Comparison of the data between untreated samples (Supplementary Figure 1C) and those treated with the lowest dose of TGF-β (5 pM) confirmed that the concentration range used here was sufficient to capture the minimal responses from low doses of TGF-β (Fig.4B, panels for 5 pM). At intermediate ligand dose (50 pM), the averaged single cell profiles showed both transient signaling (SMAD translocation) and *ctgf* responses (Fig.4B, panels for 50 pM). Above a ligand concentration of 500 pM, cells reached their maxima of SMAD nuclear shuttling and *ctgf* transcriptional response (Fig.4B, panels for 500 pM and 5 nM). *Ctgf* response levels did not appear to scale linearly with SMAD signaling, and both signals revealed the analog encoding of TGF-β dose information (Fig.4C-D). Similarly, temporal profiles of *ctgf* responses displayed ligand concentration-dependent properties (Fig.4E-F). We thus conclude that SMAD4/2 nuclear shuttling and ctgf response amplitude scale with TGF-β dosage.

**Figure 4.**
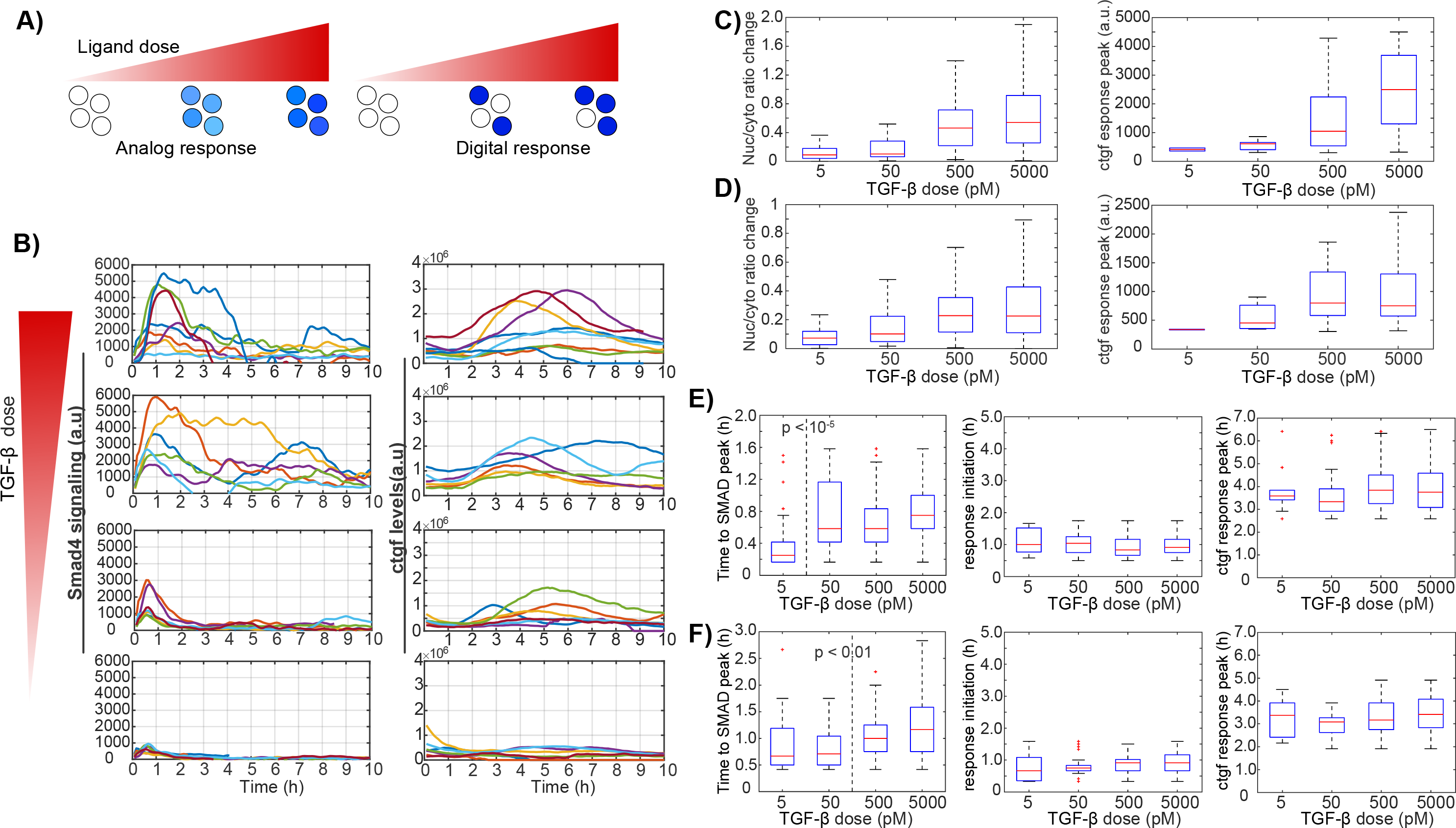
Analog encoding of TGF-β dose information by dose-dependent SMAD signaling and *ctgf* responses. A) Schematics of analog and digital responses of individual cells. White filled circles represent the unresponsive single-cells while blue shades represent the response strength. B) Stimulation dose-response of TGF-β-induced SMAD4 signaling and ctgf response in single cells. At time t=0, cells were treated with the indicated concentrations of TGF-β and representative traces for active single cells are shown. Left: traces of nuclear to cytoplasmic signal difference in SMAD4; right: *ctgf* responses in the corresponding single cells. C-D) Nuclear/cytoplasmic ratio change from stimulation to the peak and ctgf response peak upon treatment with different doses of TGF-β in the iS4 (C) and iS2 (D) cell lines. E-F) Time lag between TGF-β stimulation and peak of nuclear/cytoplasmic SMAD4/2 ratio (left), between TGF-β stimulation and *ctgf* response initiation (middle) and peak (right), in the iS4 (E) and iS2 (F) cell lines.

### *Ctgf* response dynamics are either transient or sustained

Single cells displayed a ~ 3-fold range in cell-to-cell variability of SMAD translocation and *ctgf* expression responses, both in their dynamics and amplitude. Moreover, inspection of individual cells revealed a more detailed profile of *ctgf* responses. The majority of individual *ctgf* responses showed clearly defined transients, reaching a peak after 3-4 hours, and then returning to basal levels after around 8 hours (Fig.5A). In contrast, some cells responded in a sustained manner, characterized by a weaker first response compared to the transiently activated cells, but then showed a longer lasting *ctgf* response. In this second subpopulation, cells typically also displayed a distinct second wave of response before returning to their basal values (Figure 5A), which was also less synchronous than the unique response of the transient responders. To more rigorously analyze these two types of responses, individual *ctgf* traces were categorized into two classes using k-means clustering (Fig.5B and Supplementary Figure 3A, transient - 87%, sustained - %13). Importantly, constraining the number of clusters to two allowed for a robust classification as evaluated by the silhouette score (Fig.5C), and the two identified cellular subpopulations of *ctgf* traces displayed the same distinct behavior as in our manual categorization (Figure 5D). While predominantly distinct in their dynamics, the two subpopulations also differed in their absolute levels of responses, with cells belonging to the sustained cluster displaying slightly but significantly lower initial responses.

**Figure 5.**
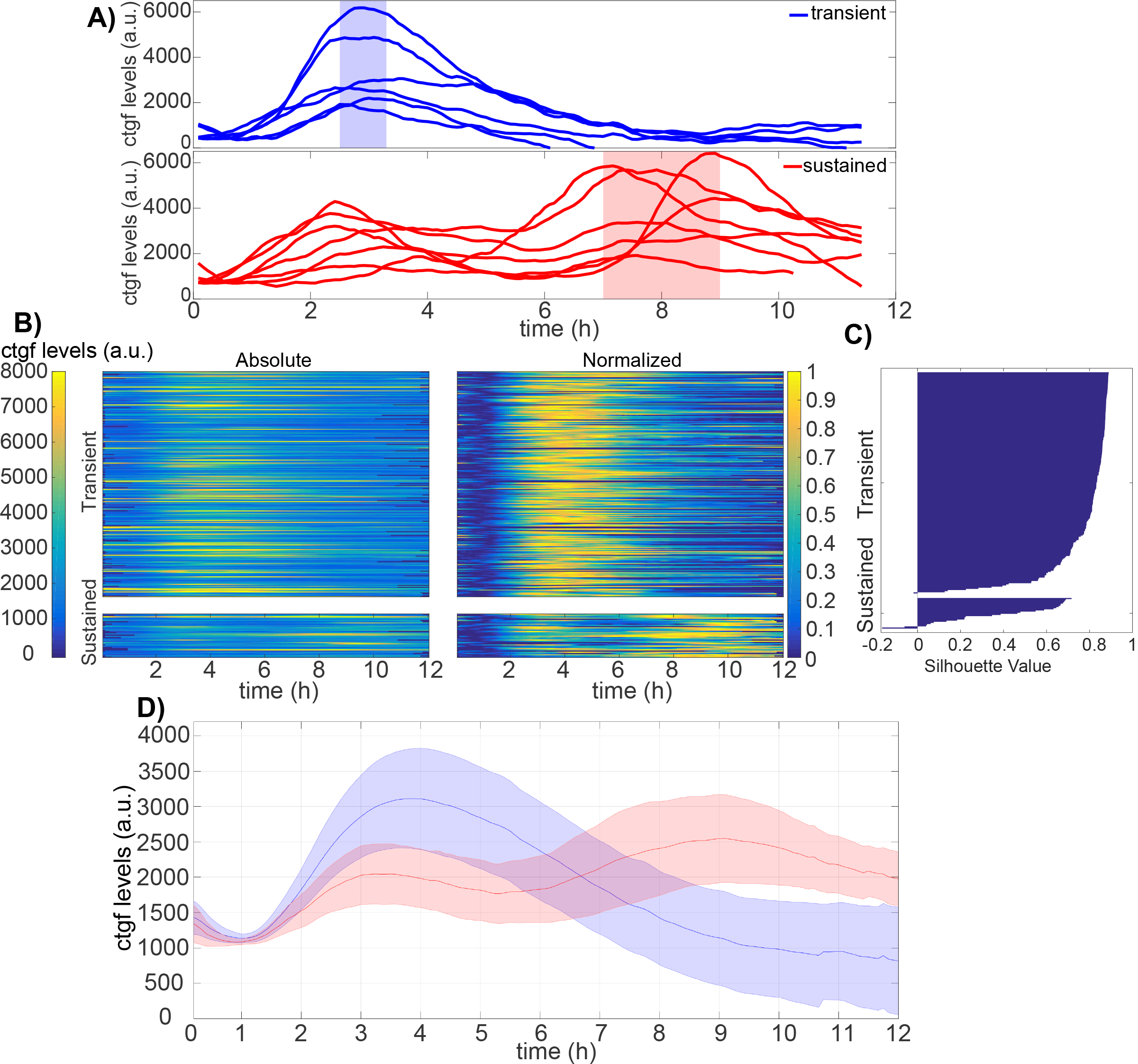
*Ctgf* response dynamics can be either transient or sustained. A) Single cell ctgf traces grouped in transient or sustained response to 5nM TGF-β stimulation with manual inspection. B) Absolute and normalized single-cell traces from TGF-β (5 nM) stimulation experiments categorized into two classes using k-means clustering (n=301 cells). C) Silhouette plot of cells sorted according to ctgf expression dynamics. D) Population-average of cells responding in a transient (blue) or sustained (red) manner. Solid lines: mean; shaded areas: SD.

### Initial SMAD4 but not SMAD2 abundance regulates the duration of the *ctgf* response

We next aimed to determine how SMAD signaling impacted on the distribution of traces in the transient or sustained classes. Interestingly, neither SMAD4 nor SMAD2 translocation dynamics displayed a significant difference in transient and sustained ctgf responses (Supplementary Figures 3B-C). In contrast, initial SMAD4 but not SMAD2 levels differed significantly between the two classes (Supplementary Figure 3D). To determine whether increased SMAD4 levels lead to a higher fraction of cells displaying a sustained *ctgf* transcriptional response, we treated the iS4 and iS2 cell lines with different doses of dox (0-200 ng/ml), and monitored the *ctgf* response after induction with 5nM of TGF-β (Fig.6). In the population-averaged data, we observed that higher SMAD4 levels resulted in a prolonged transcriptional response of *ctgf*, while increasing SMAD2 levels did not show a consistent effect (Supplementary Figure 4A). In principle, the change from a transient response at low SMAD4 concentration to a more sustained response at high SMAD4 concentration could reflect homogeneous changes of the response in the cell population, or changes in the proportion of cells responding in a transient versus sustained manner. To discriminate between these two possibilities, we employed k-means clustering of all single cell *ctgf* responses that we obtained at different dox concentrations for both iS4 and iS2 cell lines (Fig.6A-B). This resulted in two clusters, the first with transient single cell responses while the second cluster was characterized by sustained and more oscillatory target gene responses. We then analyzed the fraction of cells belonging to each cluster as a function of the dose of dox in both iS4 and iS2 cell lines. While increasing dox concentration in the iS4 cell line resulted in a higher proportion of cells responding in a sustained manner, this did not impact on the fraction of oscillating cells in the iS2 cell line (Fig.6C, Supplementary Figure 4 and Table 1). Supporting this notion, comparison of SMAD levels in the transient and the sustained classes in samples treated with 2 ng/ml and 10 ng/ml doxycycline revealed that SMAD4 levels are significantly higher in the sustained class while SMAD2 levels do not differ (Fig.6D). Therefore, the dynamics of the transcriptional response of the *ctgf* gene is influenced by the total amount of SMAD4 but not SMAD2.

**Figure 6.**
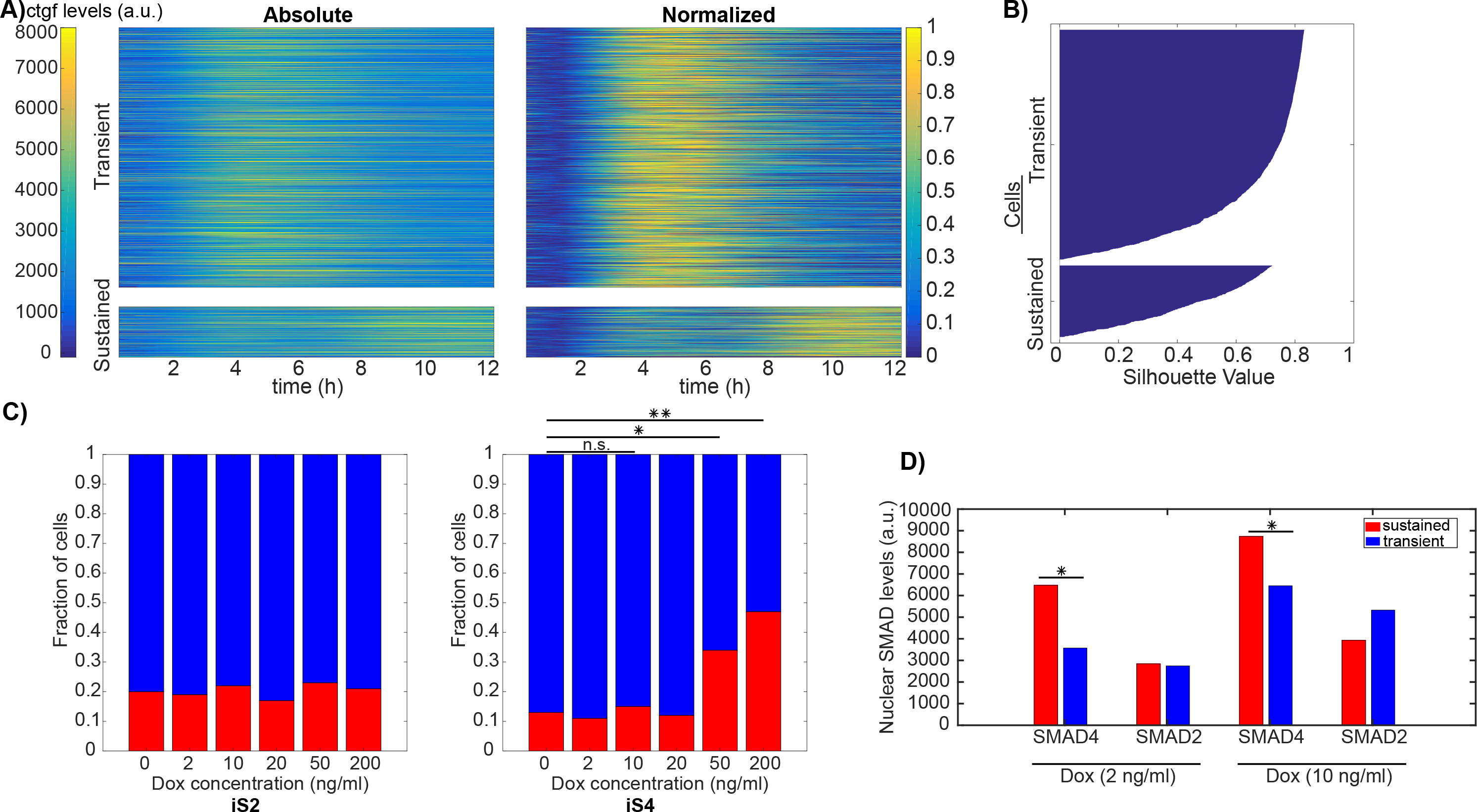
Higher SMAD4 but not SMAD2 levels increase the fraction of cells with a sustained *ctgf* response. A) Single cell *ctgf* responses after stimulation with different doses of TGF-β, clustered into transient and sustained classes using k-means clustering (n transient = 1666; n sustained = 427). Heat maps for *ctgf* traces belonging to transient (upper panel) and sustained (lower panel) are shown using absolute (left) and normalized levels of *ctgf* responses. B) Silhouette plot of cells sorted according to *ctgf* expression dynamics. C) Fraction of cells responding in a transient or sustained manner at different concentrations of dox in the iS2 (left) and iS4 (right) cell lines. (*p<0.001, **p<10e-6; Chi-squared test). D) Average SMAD4/2 expression level during the 12 hours following 5nM TGF-β stimulation, belonging to the transient and the sustained classes from samples treated with 2 ng/ml and 10 ng/ml doxycycline (*p<0.05; t-test).

**Table 1.** List of experiments performed using different doses of Dox and the number of cells classified into transient and sustained classes for each condition.

|  | iS4 |  |  |  |  |  | iS2 |  |  |  |  |  | Total |
| --- | --- | --- | --- | --- | --- | --- | --- | --- | --- | --- | --- | --- | --- |
| Dox (ng/ml) | 0 | 2 | 10 | 20 | 50 | 200 | 0 | 2 | 10 | 20 | 50 | 200 |  |
| Transient | 97 | 188 | 146 | 84 | 163 | 45 | 109 | 452 | 65 | 183 | 116 | 20 | 1668 |
| Sustained | 14 | 23 | 26 | 11 | 84 | 39 | 27 | 106 | 18 | 37 | 35 | 5 | 425 |

## Discussion

Major progress was made recently on describing dynamics of various mammalian signaling pathways ^26,27^, transcription factor nucleo-cytoplasmic shuttling ^15,28,29^, and transcription at the single cell level ^20,21,30,31^. However, there is still little known about the quantitative relationship between these parameters, which will be key to understand how cells transduce external signals into changes in gene expression level. Here we show that amplitudes of SMAD4 and SMAD2 nuclear cytoplasmic shuttling do not impact the transcriptional response amplitude of the endogenous *ctgf* gene. This suggests that this signaling pathway has evolved to allow each cell to maximize its transcriptional response once the signal has reached the transcriptional activators. In contrast, it has been shown that receptor availability is a major source of intercellular response to TGF-β signaling ^25^, and receptor endocytosis also regulates the duration of the TGF-β signaling response ^32^. We may thus speculate that TGF-β signaling is mainly regulated at the first step, *i.e.* the binding of TGF-β to its receptor, while subsequent signaling components are optimized to transfer the signal to the gene efficiently. Interestingly, we also found the temporal response of SMAD4 and SMAD2 shuttling to be fast, tightly regulated, and very quickly followed by the response in *ctgf* transcriptional activity. Therefore, the main temporal limitation to the response lies within the accumulation of gene expression products, which depends on the half-lives of the mRNA and proteins produced. Interestingly, both *ctgf* mRNA and proteins are short-lived ^33^, which should allow very rapid maximal expression of the CTGF protein in response to TGF-β signaling. Continuous TGF-β stimulation has been described as generating transient responses, while consecutive pulse stimulations was shown to result in sustained activation ^15,17^. This heterogeneity in the response profile of individual cells could be due to differences in negative feedback efficiency ^16,27,34^ or secondary mechanisms of SMADs recruiting activators and suppressors, thereby shaping the transcriptional response later after stimulation ^35^. Surprisingly, we found that SMAD4 but not SMAD2 expression levels regulate the probability of cells to display a transient versus a sustained *ctgf* transcriptional response. While the mechanistic basis underlying this observation remains unclear, it is possible that higher SMAD4 levels allow to overcome the negative feedback generated after TGF-β stimulation.

Together with TGF-β, CTGF participates in wound healing to reconstitute a properly arranged connective tissue ^19^. In normal adult fibroblasts, TGF-β controls the expression of *ctgf* which induces fibroblast proliferation and production of extracellular matrix ^36,37^. However, uncontrolled CTGF expression is generally associated with pathological forms of fibrosis characterized by uncontrolled scarring ^38–40^ or certain types of cancer ^41^. Due to its angiogenic functionality, high basal levels of CTGF can provide favorable environments for metastasis when induced by the TGF-β pathway ^42–44^. It was shown that depletion of SMAD4 in these cells causes a substantial reduction on metastatic potential ^45^. While the kinetic profile of *ctgf* expression in these contexts is unknown, our observations suggest that higher SMAD4 levels may also allow more sustained *ctgf* transcription activity in the context of fibrosis and tumor metastasis. Further studies shall address the mechanistic basis of how elevated SMAD4 levels generate sustained *ctgf* transcriptional activity.

## Methods

### Construction of lentiviral plasmid constructs

NLuc was amplified from a synthetic construct using primers 5’-CGT AAA ACC GGT CGA ATG GTC TTC ACA CTC GAA G-3’ and 5’-AGA CAT ATT GTC CAT GTC GAC CGC CAG AAT GCG-3’. Smad4 was amplified from cDNA synthesized from NIH-3T3 RNA using primers 5’-ATG GAC AAT ATG TCT ATA ACA A-3’ and 5’-CGA ACA CGT GGT CGA TCA GTC TAA AGG CTG TGG G-3’. pLVTRE3G-NLuc-Smad4 was constructed by three-fragment In-fusion (Clontech) cloning of pLVTRE3GMCS ^46^ digested with SalI, NLuc, and Smad4. Smad2 was amplified from cDNA synthesized from NIH-3T3 RNA using primers 5’-CAT GTC GAC ATG TCG TCC ATC TTG CCA TT-3’ and 5’-CAT CAT ATG TTA CGA CAT GCT TGA GCA TCG-3’ and ligated into pLVTRE3G-NLuc-Smad4 digested with SalI and NdeI (NEB) using T4 DNA ligase (NEB). pLV-PGK-rtTA3G-IREShygro was constructed as described previously ^23^. All constructs were verified by Sanger sequencing.

### Lentiviral vector production and generation of stable cell lines

Lentiviral vector production was performed by co-transfection of HEK 293T cells with the lentiviral construct, the envelope (PAX2) and packaging (MD2G) constructs using calcium phosphate, and concentrated 120-fold by ultracentrifugation as described previously ^20^. NIH-3T3 GT:ctgf cells ^20^ were transduced with 120-fold concentrated virus carrying pLV-PGK-rtTA3G-IREShygro followed by selection with 200 μg/ml Hygromycin. Subsequently, these cells were infected with either pLVTRE3G-NLuc-SMAD4 or pLVTRE3G-NLuc-SMAD2, followed by selection with 2 μg/ml Puromycin. These two stable cell lines (ctgf-SMAD2 and ctgf-SMAD4) were seeded at clonal density and clones were picked manually to obtain more homogeneous expression levels of the transgene.

### Cell culture

The GT:ctgf, iS4, iS2 NIH-3T3 cell lines and HEK 293T cells (ATCC) were cultured in DMEM (Thermofisher; 41966029), supplemented with 10% fetal bovine serum (Thermofisher, 10270106) and 1% penicillin/streptomycin (BioConcept, 4-01F00H), at 37°C and 5% CO_2_. Cells were grown in 100 mm cell culture plates up to a confluence of 70% and split 1/6 every 2-3 days.

### Single-cell luminescence time-lapse microscopy

Luminescence time-lapse recordings were performed on an Olympus LuminoView LV200 microscope equipped with an EM-CCD cooled camera (Hamamatsu photonics, EM-CCD C9100-13), a 60x magnification objective (Olympus UPlanSApo 60x, NA 1.35, oil immersion) in controlled environment conditions (37°C, 5% CO2). To discriminate the luminescence signals from Nluc and Fluc, 700nm LP filter (Chroma) for Fluc and 460/36nm band-pass filter (Chroma) for Nluc imaging were used. One day before the experiment, cells were seeded on 35mm fluorodishes (WPI Inc, FD35-100). Before imaging, the medium was supplemented with 500μM Luciferin (NanoLight Technology 306A) and 0.5μl of RealTime Glo Cell Viability Assay Substrate (Promega G9711). Images were acquired every 3 minutes in the Nluc channel and every 2 minutes in the Fluc channel with a cycle time of 5 minutes up to 24 hours. Cells were recorded for 0-6 hours before stimulation with mouse TGF-β1 (eBioscience, 14-8342-62).

Analyses of intensities from the two channels revealed no detectable bleed-through signal observed in 460/36nm (Nluc) channel due to luminescence emitted by Luciferin while 1.5% of the Nluc signal was visible in the Fluc channel. This fraction of Nluc signal was subtracted from Fluc signal measured in single cells when analyzing dual-luminescence time-lapse imaging data.

### Cell tracking

Tracking of cells was performed using CAST (Cell Automated Segmentation and Tracking platform^47^ (Cell Automated Segmentation and Tracking platform). After preprocessing, images are convolved with a family of cell-like filters and converted to binary images using an adaptive threshold to define nuclear regions. The cytoplasmic region was defined using an annulus around the nucleus. To remove spurious detections of cells, an additional optional step was utilized to filter out short trajectories before the gap closing, merging and splitting steps, thus preventing these from being linked together into a spurious trajectory.

### Western Blotting

Cells were grown to reach confluency before performing protein extraction. For time-lapse TGF-β stimulation samples, cells were stimulated 1 day after seeding and collected with counting at indicated time points by trypsinization and centrifugation. Cells were then lysed in RIPA buffer (50 mM Tris pH 7.4, %1 NP-40, 0.5% NaDeoxycholate, 0.1% SDS, 150 mM NaCl, 2 mM EDTA), supplemented with 1mM PMSF (AppliChem A0999.0005) and Protease Inhibitors (Sigma P8340- 5ML). Samples were left on ice for 30 minutes and then spun down at 14000g for 15 minutes at 4°C. The protein concentration of the supernatant was determined by performing a Bicinchoninic acid assay (BCA) (ThermoFisher 23235) and 15μg of protein were mixed with Laemmli sample buffer (Invitrogen NP0007) and loaded on an SDS gel (BioRad 456-1094) for separation (SDS Running Buffer 25mM Tris, 190mM Glycine, 0.1%SDS). Proteins were subsequently transferred from the gel onto a nitrocellulose membrane using a dry transfer system (Merck IB21001, iBlot 2 Dry Blotting System). Antibodies against C-terminally phosphorylated Smad2 (3108; Cell Signaling Technologies) (dilution 1:1,000), Smad2/3 (610482; BD Transduction Labs) (dilution 1:1,000), and Smad4 (B-8; Santa Cruz) (dilution 1:500) were used. The membrane was blocked with 5% bovine serum albumin or 5% milk (Roth T145.3) in TBS-T (for phosphorylated samples) or PBS-T followed by incubation with primary antibody overnight. The following primary antibodies with given dilutions were used; Smad4 (ABE21; Merck, dilution 1:2000), pSmad2 (D27F4; Cell Signaling Technologies, dilution 1:1000), Smad2 (D43B4; Cell Signaling Technologies, dilution 1:1000). Membranes were subsequently washed shortly and incubated with HRP-conjugated secondary antibodies in %5 nonfat dry milk in TBS-T (for phosphorylated samples) or PBS-T. The following secondary antibodies with given dilutions were used; anti-mouse IgG-HRP (W402B; Promega, dilution 1:10 000) and anti-rabbit IgG-HRP (W401B; Promega, dilution 1:10 000). Membranes were then washed in TBS-T/ PBS-T and imaged. Protein bands were visualized using Clarity Western ECL Substrate (BioRad 170-5060). Images were captured using a Vilber-Fusion chemiluminescence system (Molecular Imaging Vilber Fusion FX7) and analyzed using ImageJ.

### Immunofluorescence

NIH-3T3 cells were fixed for 15 min with ice-cold 4% PFA (AppliChem A0877,0500) in PBS, permeabilized and blocked with chilled PBS-Triton (AppliChem A1388,0500) and 1% FBS for 30 - 60 min. Samples were incubated with the primary antibody in PBS and 1% FBS overnight at 4°C, washed twice in PBS, and incubated with the secondary antibody in PBS and 1% FBS for 45 - 60 min. Samples were then washed three times with 0.1% PBS-Tween (Fisher Scientific BP337- 500), incubated with 1 μg/mL DAPI for 15 minutes, washed twice with 0.1% PBS-Tween and once with PBS.

To quantify signals in immunofluorescence samples, a semi-automated image analysis pipeline built in the Cell Profiler software (www.cellprofiler.org) was used. DAPI staining was used to precisely locate nuclear regions and SMAD4 staining was used to define cellular borders and to determine cytoplasmic regions. Manual correction was performed for erroneous detections. Based on defined nuclear and cytoplasmic regions, signal intensities from samples were extracted. Background subtraction was performed using control samples.

### Clustering analysis

Clustering was performed using an unsupervised *k-means* algorithm provided in MATLAB (MathWorks) software package (*k-mean*, Lloyd, 1982). The function uses a modified version for initialization ^49^. Before the algorithm was executed, a heuristic choice of two clusters was made to comply with the manual inspection of single cell traces and the identification of two distinct behaviors. For the k-means clustering, we used the correlation matrix as the distance metric to determine subgroups with similar dynamic patterns.

Silhouette plots provide a graphical representation of how well each member belonging to a cluster corresponded to other members in the same cluster rather than the other cluster ^50^. The silhouette value for the *i*-th cell, *S*_*i*_, is

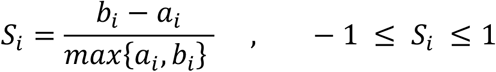

where *a*_*i*_ represents the average distance from the *i*-th point to the other points in the same cluster as *i*, and *b*_*i*_ is the minimum average distance from the *i*_th_ point to points in a different cluster, minimized over clusters. The silhouette score ranges from −1 to +1, where a high value indicates that the object is well matched to its own cluster and poorly matched to neighboring clusters.

## Supporting information

## Acknowledgments

O.T. is funded by a SystemsX.ch IPhD grant 51PH_0125979 to David Suter and Felix Naef.

## Author Contributions

Conceptualization, O.T., F.N. and D.M.S.; Methodology, O.T. and E.T.F; Software, O.T. and F.N; Formal Analysis, O.T., D.M.S. and F.N; Investigation, O.T., E.T.F, F.N. and D.M.S.; Resources, D. M.S. and .N., Writing – Original Draft, O.T., F.N and D.M.S.; Writing – Review & Editing, O.T., E. T.F, F.N. and D.M.S.; Funding Acquisition, F.N. and D.M.S; Supervision, F.N. and D.M.S.

## Data availability

The datasets generated during the current study are available from the corresponding author on reasonable request.

