## Supplementary material for "Quantitative relationships between SMAD dynamics and target gene activation kinetics in single live cells"

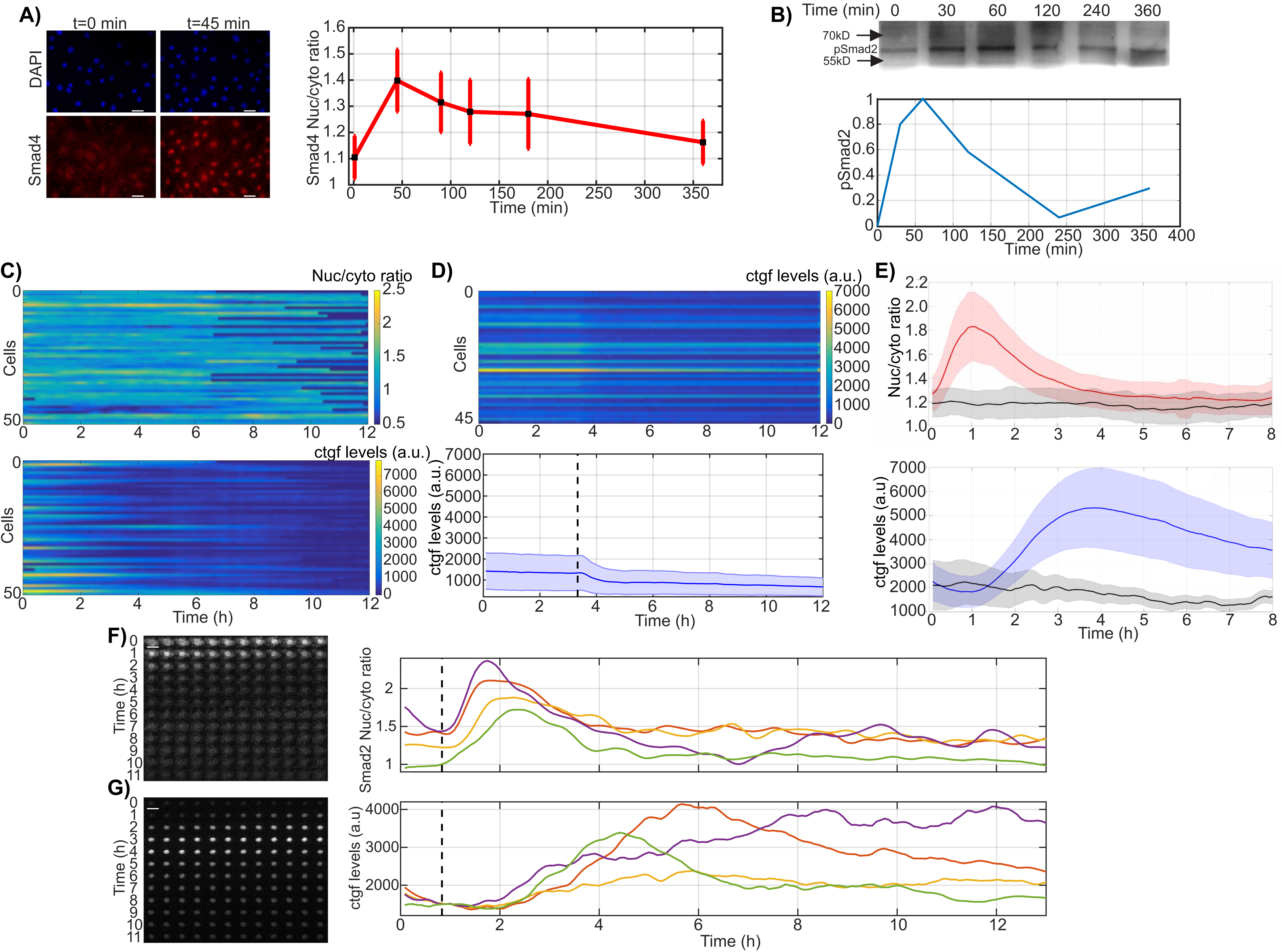

A)
t=0 min
t=45 min
DAPI
Smad4
Smad4 Nuc/cyto ratio
Time (min)
B)
Time (min) 0 30 60 120 240 360
70kD
pSmad2
55kD
pSmad2
Time (min)
C)
Nuc/cyto ratio
Cells
ctgf levels (a.u.)
Cells
Time (h)
D)
ctgf levels (a.u.)
Cells
ctgf levels (a.u.)
Time (h)
E)
Nuc/cyto ratio
ctgf levels (a.u)
Time (h)
F)
Time (h)
Smad2 Nuc/cyto ratio
G)
Time (h)
ctgf levels (a.u)
Time (h)

**Supplementary Figure 1. Localization and dynamics of SMAD2/4 and *ctgf* response.**

A) Time course immunofluorescence analysis of SMAD4 translocation in response to 5nM TGF-β stimulation.

B) Western blots of pSMAD2 in total cell lysates after 5nM TGF-β stimulation. The quantification shows normalized band intensity with min-max scaling. The cropped blot is used in the figure and the full length Western blot scans for the cropped images are shown in Supplementary Figure 5.

C) SMAD4 localization changes (top) and *ctgf* expression (bottom) without stimulation.

D) Individual cell traces (top) and population average (bottom) of *ctgf* expression levels in samples treated with the TGFβ-receptor antagonist SB-431542 (dashed black line). Solid line: mean; shaded areas: SD.

E) Comparison of SMAD4 translocation (top) and *ctgf* expression (bottom) in unstimulated cells (n= 58, gray) or cells stimulated with 5nM TGF-β (red and blue). Solid line: mean; shaded areas: SD. Scale bar: 20 µm.

F-G) Time series images of a tracked single cell of iS2 cell line on both Nluc (left) and Fluc (right) channel by luminescence microscopy. Time resolution: 5 minutes. Four individual traces are shown for both SMAD2 translocation and *ctgf* response.

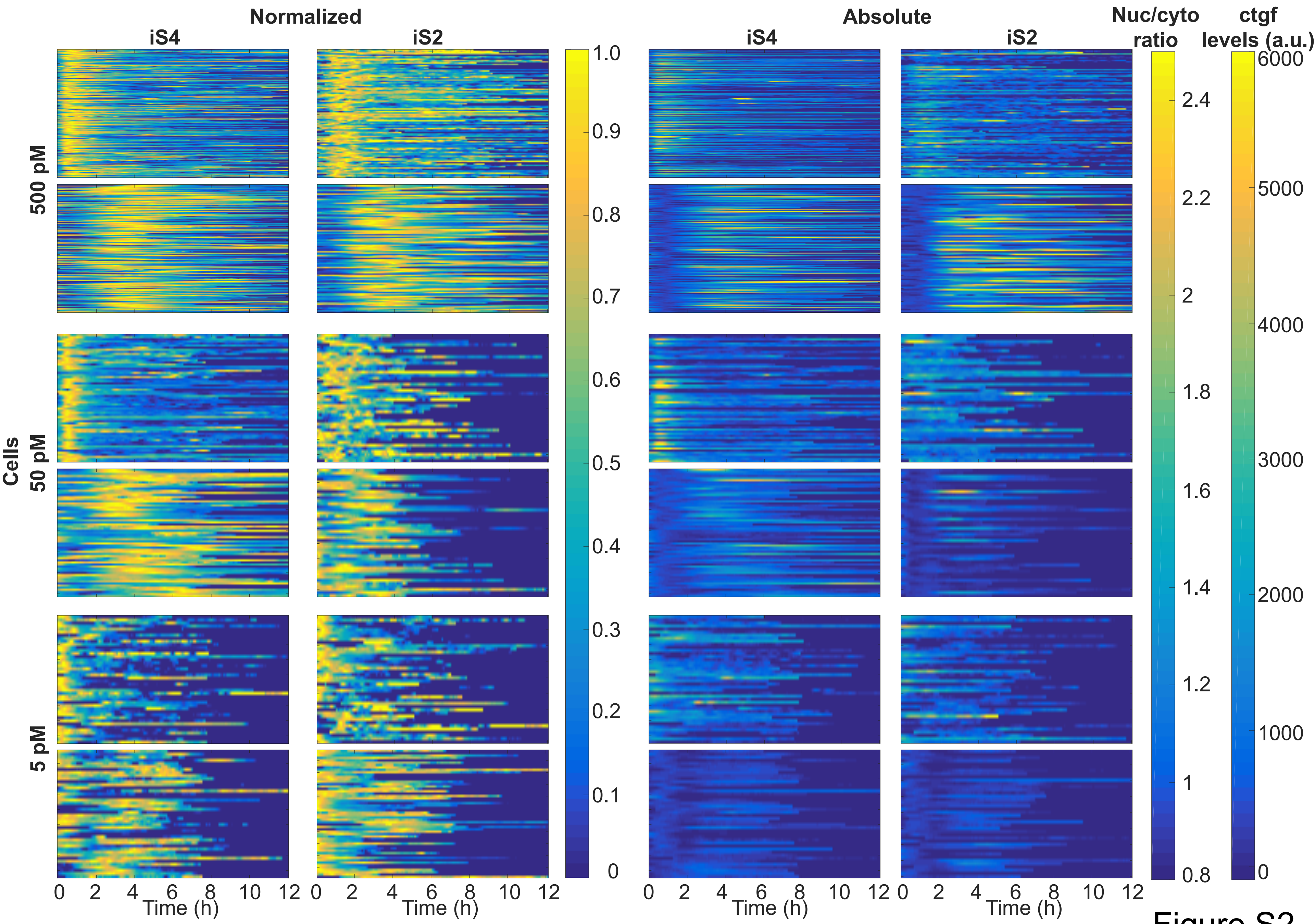

Normalized
Absolute
iS4
iS2
iS4
iS2
Nuc/cyto ratio
ctgf levels (a.u.)
Cells
500 pM
50 pM
5 pM
Time (h)
Time (h)
Time (h)
Time (h)


Figure S2

**Supplementary Figure 2. Heat maps of for single cell traces of SMAD4/2 translocation and *ctgf* responses to TGF-β stimulation.**

Cells were treated with different doses of TGF-β (500-50-5 pM) at time t=0 and response traces for all cells are shown both in normalized (left) and absolute (right) terms. For each experiment, SMAD translocation (top) and ctgf response (bottom) are shown.

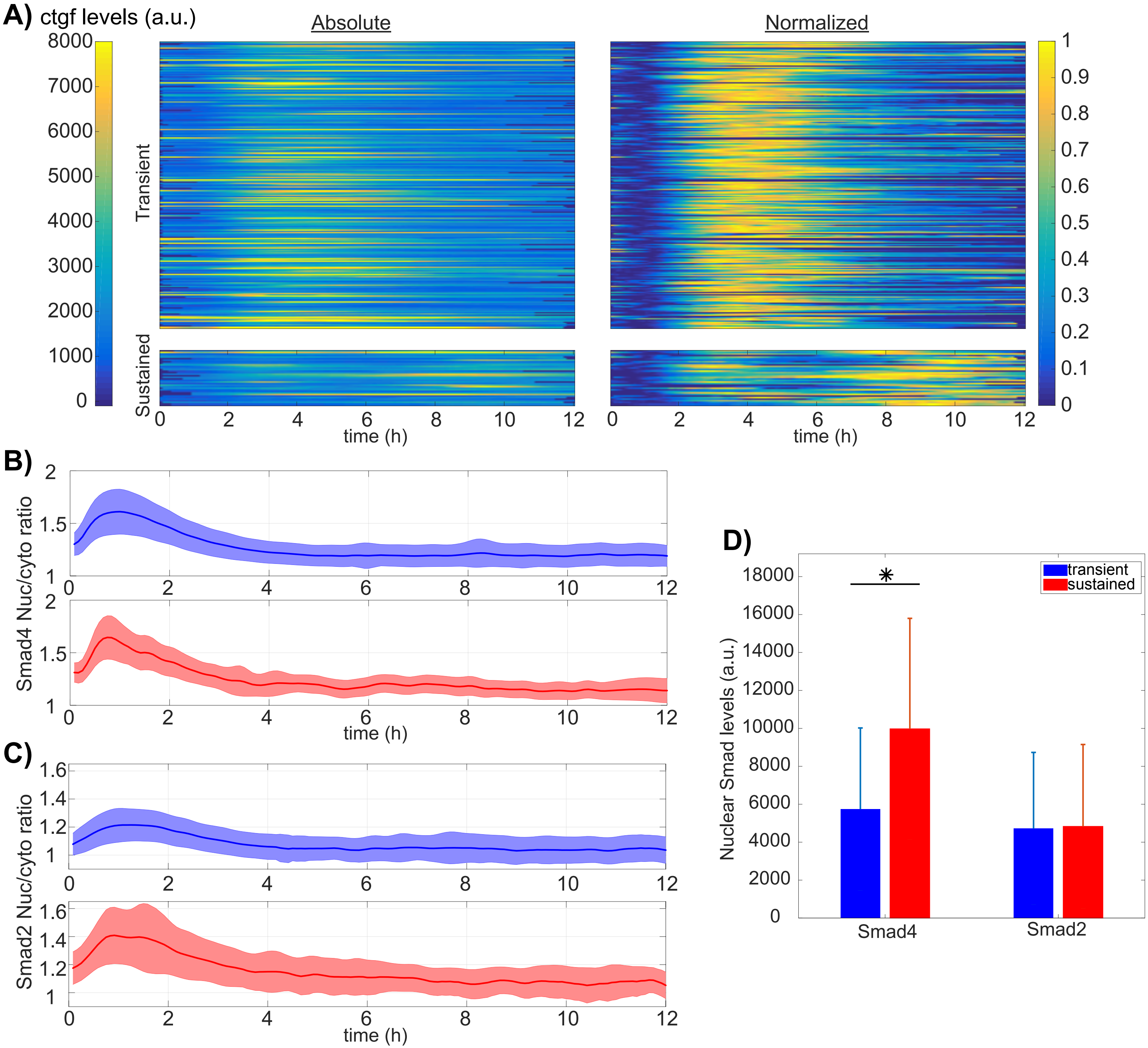
A) ctgf levels (a.u.)
Absolute
Normalized
Transient
Sustained
time (h)
B)
Smad4 Nuc/cyto ratio
time (h)
C)
Smad2 Nuc/cyto ratio
time (h)
D)
Nuclear Smad levels (a.u.)
transient
sustained
*
Smad4
Smad2

Figure S3

**Supplementary Figure 3. SMAD shuttling dynamics and levels in cells responding in a transient and sustained manner TGF-β stimulation.**

A) Decomposition of single cell *ctgf* responses into transient and sustained classes using k-means clustering in iS2 cell line upon 2ng/ml dox treatment and 5nM TGF-β stimulation. Heatmaps for ctgf traces belonging to transient (upper panels) and sustained (lower panels) are shown, quantified as both absolute (left) and normalized (right) levels (min-max scaled, 0 to 1).

B-C) Average SMAD nuclear to cytoplasmic ratio trajectories belonging to two classes identified with clustering (blue-transient, red-sustained ), in iS4 (B) and iS2 (C) cell lines.

D) Average SMAD4/2 expression level during the 12 hours following 5nM TGF-β stimulation, belonging to the transient and the sustained classes from samples treated with all doses of dox. Error bars: SD; *p<0.005 (t-test).

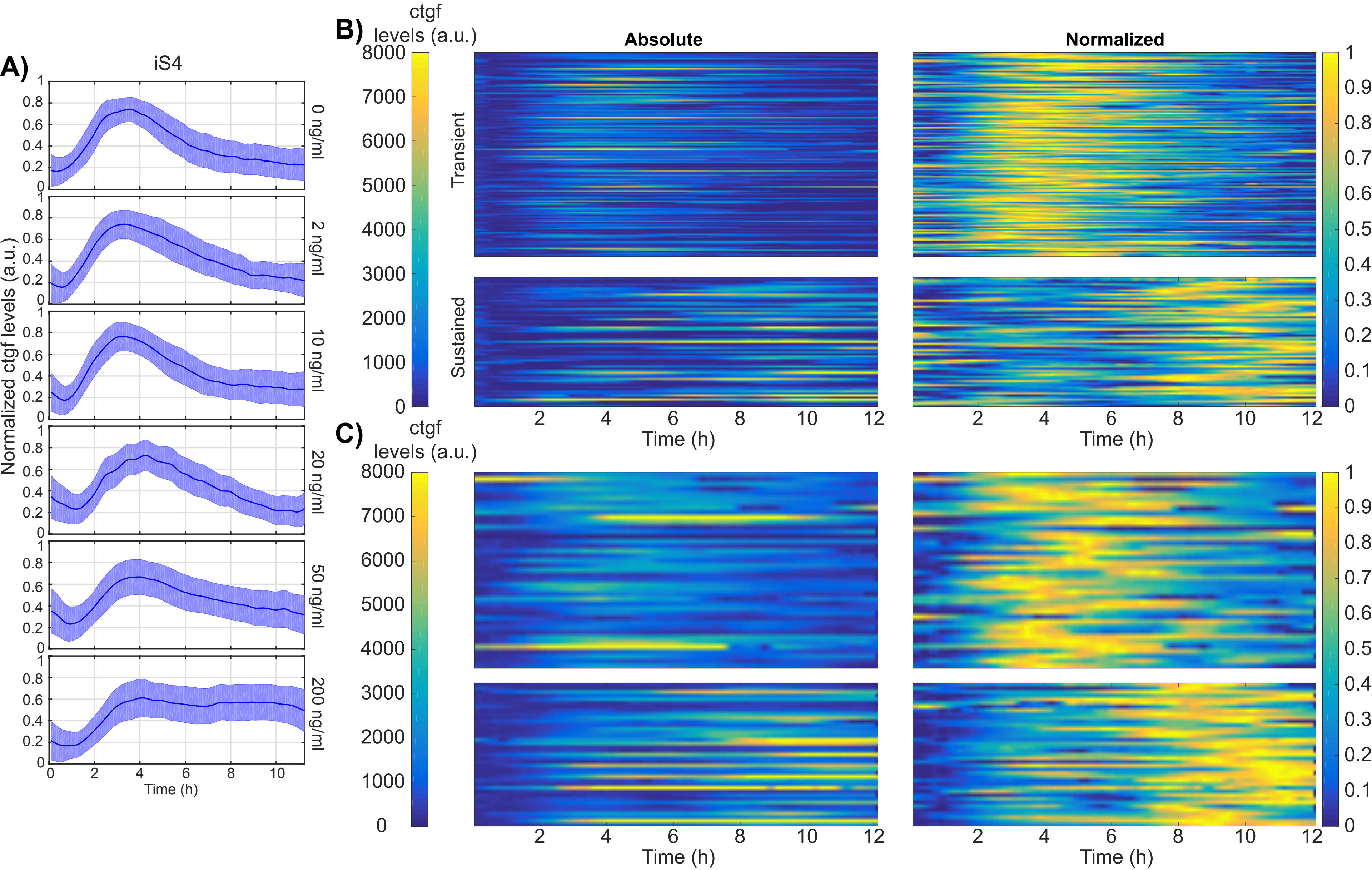

A)
iS4
Normalized ctgf levels (a.u.)
0 ng/ml
2 ng/ml
10 ng/ml
20 ng/ml
50 ng/ml
200 ng/ml
Time (h)
B)
ctgf levels (a.u.)
Absolute
Normalized
Transient
Sustained
Time (h)
C)
ctgf levels (a.u.)
Time (h)


Figure S4

**Supplementary Figure 4. TGF-β-induced SMAD signaling and *ctgf* response in iS4 cell line treated with different doses of Dox.**

A) Population averages of *ctgf* responses upon 5nM TGF-β stimulation in samples treated with varying doses of doxycycline in the iS4 cell line.

B-C) Clustering of single cell *ctgf* responses into transient and sustained classes in iS4 cell line treated with 50ng/ml (B) and 200ng/ml (C) of dox.

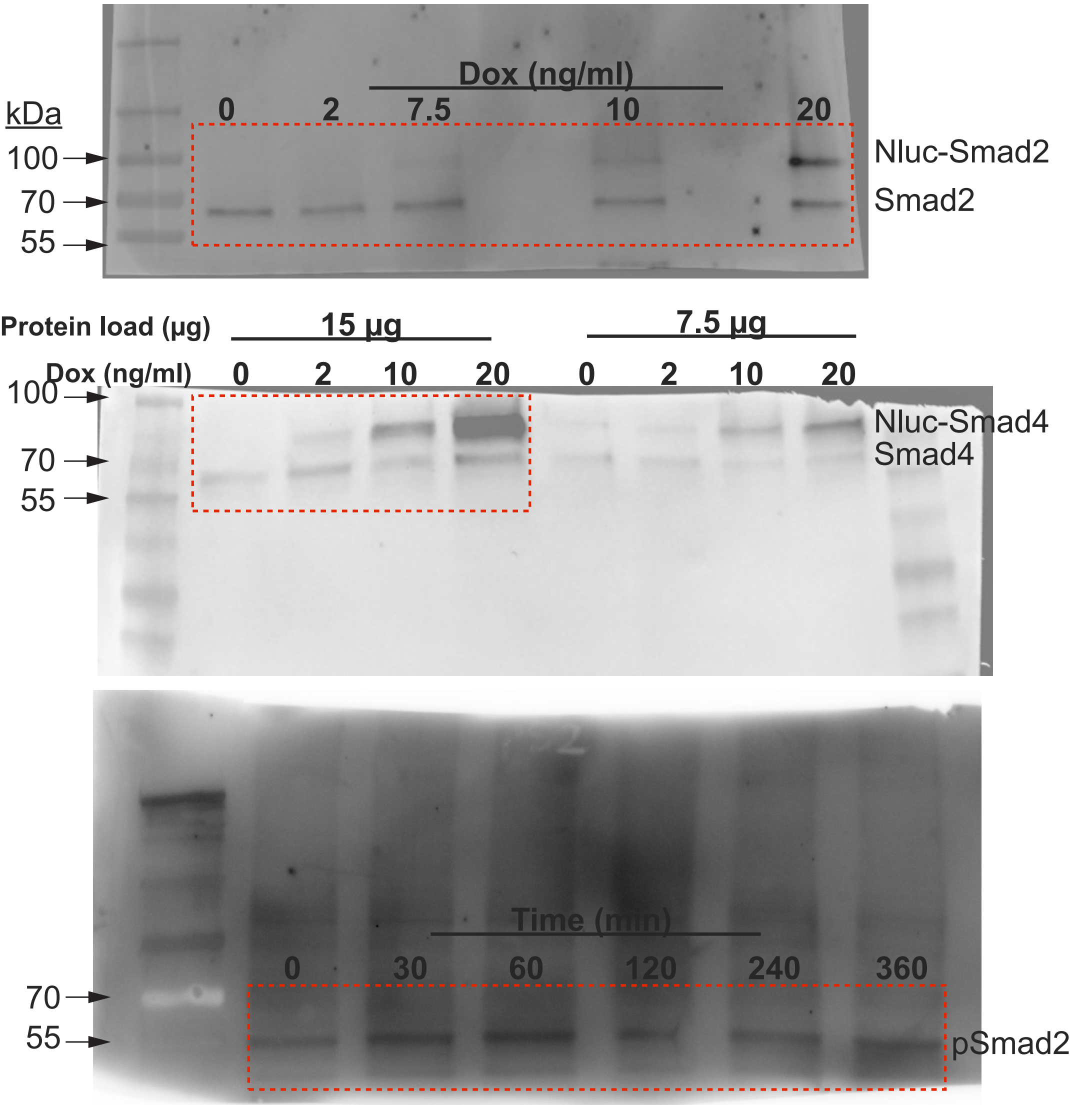

kDa
100
70
55
Dox (ng/ml)
0
2
7.5
10
20
Nluc-Smad2
Smad2
Protein load (µg)
15 µg
7.5 µg
Dox (ng/ml)
0
2
10
20
0
2
10
20
100
70
55
Nluc-Smad4
Smad4
Time (min)
0
30
60
120
240
360
70
55
pSmad2

**Figure S5. Full images of Western blots shown in Figure 1 and Supplementary Figure 1.** Red dotted lines indicate the cropping locations. Brightness was adjusted during processing of these blots.
